# Spectacle: Faster and more accurate chromatin state annotation using spectral learning

**DOI:** 10.1101/002725

**Authors:** Jimin Song, Kevin C. Chen

## Abstract

Recently, a wealth of epigenomic data has been generated by biochemical assays and next-generation sequencing (NGS) technologies. In particular, histone modification data generated by the ENCODE project and other large-scale projects show specific patterns associated with regulatory elements in the human genome. It is important to build a unified statistical model to decipher the patterns of multiple histone modifications in a cell type to annotate chromatin states such as transcription start sites, enhancers and transcribed regions rather than to map histone modifications individually to regulatory elements.

Several genome-wide statistical models have been developed based on hidden Markov models (HMMs). These methods typically use the Expectation-Maximization (EM) algorithm to estimate the parameters of the model. Here we used spectral learning, a state-of-the-art parameter estimation algorithm in machine learning. We found that spectral learning plus a few (up to five) iterations of local optimization of the likelihood outperforms the standard EM algorithm. We also evaluated our software implementation called **Spectacle** on independent biological datasets and found that **Spectacle** annotated experimentally defined functional elements such as enhancers significantly better than a previous state-of-the-art method.

**Spectacle** can be downloaded from https://github.com/jiminsong/Spectacle.

## 1 Introduction

Identifying regulatory elements in the genome is a challenging problem with applications in understanding many aspects of biology, including the molecular mechanisms of disease. In particular, cell-type specific gene regulation cannot be explained solely by genome sequence data because the genome is essentially the same in most cell-types. Recently, the ENCODE project (ENCODE Project Consortium 2012) produced a wealth of epigenetic data using biochemical assays and next-generation sequencing (NGS) technologies. This data has the potential to significantly improve our understanding of cell-type specific gene regulation. In addition to ENCODE, the International Human Epigenome Consortium (IHEC, http://ihec-epigenomes.org/) also aims to produce reference maps of 1000 human epigenomes. It includes several major projects such as BLUEPRINT and the Roadmap Epigenomics Project (Adams *et al.* 2012; Bernstein *et al.* 2010). Here we refer to epigenetic data as histone modification, DNA methylation, transcription factor binding, open chromatin and other related types of data generated by biochemical assays. In particular, histone modification refers to post-translational modification of histones where chemical groups such as methyl, acetyl, phosphate, ubiquitin, SUMO, ADP-ribose etc are added or deleted to histone tails (Portela and Esteller 2010). The “epigenome” provides another dimension to our understanding of the human genome and it can be used to identify cell-type specific gene regulatory elements or disease variants (Rivera and Ren 2013). Genomic features delineated by epigenomic marks have been linked to genome-wide association study (GWAS) loci (Ernst *et al.* 2011; Maurano *et al.* 2012). Since up to 93% of GWAS SNPs may be located in non-coding regions, such results give hope that one might be able to fine-map the causal disease variants.

Several systematic methods have been developed to identify regulatory elements or annotate chromatin states from epigenetic data (mainly, histone modification data) not only in the human genome but also in the Drosophila, mouse and yeast genomes (Ernst and Kellis 2010; Filion *et al.* 2010; Kharchenko *et al.* 2011; Hoffman *et al.* 2012; Shen *et al.* 2012; Wang *et al.* 2012; Biesinger *et al.* 2013; Hoffman *et al.* 2013; Lai and Buck 2013; Won *et al.* 2013; Zeng *et al.* 2013). Among these methods, Hidden Markov Models (HMMs) were repeatedly used as the underlying probabilistic model. HMMs (Hidden Markov Models) are widely used in many different fields including natural language processing and computational biology. In order to annotate chromatin states, we can design an HMM composed of hidden states which represent chromatin states such as transcription start sites (TSSs), promoters, enhancers, transcribed regions, non-coding RNA regions etc., and emission states which represent combinations of histone marks.

To estimate the parameters of an HMM in an unsupervised way (i.e. without access to labelled examples), the EM (Expectation-Maximization) algorithm has been the standard algorithm for a long time (Dempster *et al.* 1977; Rabiner 1989). The EM algorithm is a likelihood approach that iteratively converges to a local optimum in the likelihood of the data. It is iterative so it is often slow and it is somewhat arbitrary to decide when to stop the iterations. Since EM is not guaranteed to find a global optimum, effective parameter initialization is important to achieve good practical performance (Huang *et al.* 2001).

Here we employed spectral learning, a state-of-the-art algorithm in machine learning to estimate the parameters of an HMM, which was recently developed by (Hsu *et al.* 2012). It is a Method of Moments approach which predates the Maximum Likelihood approach (Pearson 1895) and is different by nature from the EM algorithm. Instead of attempting to find the maximum likelihood solution, it expresses various unobservable moments of the model as functions of the parameters, sets these moments equal to the sample moments estimated from the data and solves these equations to estimate the parameters.

One advantage of spectral learning is that it is provably correct under mild assumptions, specifically that the transition and emission matrices of the HMM are full rank and that the initial state vector is strictly greater than 0 in all coordinates. It has theoretically provable polynomial sample complexity bounds, which gives us a way to analyze the minimum sample size needed to infer the HMM parameters correctly. It does not suffer from local optima issues so it does not depend on the parameter initialization.

For practical applications, the spectral algorithm can be much faster than the EM algorithm. The main computation in spectral learning is to compute the SVD (singular value decomposition) of a matrix whose dimension depends on the number of histone marks. Thus the time complexity of the computation is not dominated by the genome size or the number of hidden states.

Our main technical contribution is to develop a practical implementation of spectral learning for HMMs for annotating chromatin domains that does not suffer from numerical instability issues. The original method of (Hsu *et al.* 2012) learned the HMM parameters only up to an unknown invertible transformation. The transformed parameters are still useful for computing the likelihood of the data and predicting future observations but they cannot be used to annotate the hidden chromatin states. Mossel and Roch showed how to infer the HMM parameters explicitly for the special case where the state and observation spaces have the same dimension (Mossel and Roch 2006) and (Hsu *et al.* 2012) extended their method to more general HMMs. However, the published method can be unstable if the eigenvalues corresponding to the chromatin states are not well separable and the method relies on injecting noise into the eigenvector computation to spread the eigenvalues (Hsu *et al.* 2012). We have improved the method in a deterministic and principled way to infer the (untransformed) HMM parameters by using the empirical observation that for chromatin data, the state space is much smaller than the observation space and there is a major observation that we call an “anchor observation” for each state.

In addition, the method of (Hsu *et al.* 2012) assumed that we have access to many short, independent samples from the HMM. We have modified their method to our application where we have a few long samples from the HMM by rewriting the formula for the transition matrix parameters in a form that does not depend on the initial state probability distribution. Finally, we have contributed empirically developed solutions for several technical issues for our application of chromatin state annotation in order to deal with noise in the data, overfitting and negative probability values.

We ran our method called **Spectacle** (SPECTral learning for Annotating Chromatin Labels and Epigenomics) for different numbers of states and multiple cell types. In most cases, **Spectacle** found a higher likelihood of the data than ChromHMM (Ernst and Kellis 2012) which has the same underlying statistical model but uses the EM algorithm for inference. Moreover, **Spectacle** was consistently much faster than ChromHMM for training parameters. Since ENCODE generated data for 147 cell types and much more epigenomic map data is expected to be produced in the future, we believe that **Spectacle** will scale up much better to tackling such data sets. In addition we picked one first tier cell type from ENCODE, GM12878, to perform external validations. We utilized independent biological datasets such as TSS, transcribed regions, lincRNAs (long intergenic non-coding RNAs) and enhancers to evaluate our method. Overall, our method uncovered patterns of histone modification marks associated with biological datasets significantly better than ChromHMM. Finally, **Spectacle** was implemented in Java by modifying the ChromHMM code. **Spectacle** should be easily portable to most desktops and the source code is freely available online.

## 2 Methods

### 2.1 Related work

To identify regulatory elements in the human genome from histone modification data, there are two broad classes of approaches-supervised and unsupervised learning. Supervised learning associates patterns of histone modifications with known chromatin states such as enhancers, promoters and non-coding RNAs. It can have good performance for known chromatin states but it requires the availability of known examples and it cannot discover new types of chromatin states (Guttman *et al.* 2009; Heintzman *et al.* 2009; Filion *et al.* 2010; Yip *et al.* 2012; Won *et al.* 2013). Unsupervised learning discovers patterns of histone modifications for each chromatin state *de novo* (Lian *et al.* 2008; Jaschek and Tanay 2009; Ernst and Kellis 2010; Ucar *et al.* 2011; Ernst and Kellis 2012; Hoffman *et al.* 2012).

ChromHMM (Ernst and Kellis 2010, 2012) uses a multivariate Hidden Markov Model (HMM) to predict chromatin states from histone mark data. Each chromosome is segmented into non-overlapping regions of 200 bp (base pair) and each segment has a binary value representing the presence/absence of each histone mark. Given a fixed number of hidden states, each segment emits a specific combination of histone marks in a hidden state. To perform unsupervised learning of the HMM parameters, they used the standard EM algorithm (Dempster *et al.* 1977), also called the Baum-Welch algorithm in the context of HMMs. For the EM algorithm to avoid local optima, it is important to initialize the parameters well. Instead of initializing the parameters at random, they proposed a heuristic method called “information” (Ernst and Kellis 2012) which we briefly describe in the Supplementary Material. ChromHMM ran the EM algorithm to convergence, that is, the difference between the likelihood of the current iteration and that of the previous iteration was less than 0.001. Regardless of convergence, the maximum number of iterations was set to 200.

Hoffman *et al.* 2012 used a generalization of HMMs called “dynamic Bayesian networks” (DBN) to model chromatin states. For example, their program, Segway, learns how long each state lasts by adding another hidden state called a *countdown* variable. Segway has high-resolution since it uses a segment size of 1 bp. However, it is much slower than ChromHMM because of the high-resolution and the complexity of the model. Nevertheless, the performance of Segway on biological data sets seems to be similar to ChromHMM (Hoffman *et al.* 2013). Since the state space is much bigger than ChromHMM, it might be harder to find the global optimum of the likelihood. Currently, Segway cannot be run on a desktop but needs to be run on a compute cluster. Thus we did not use it for comparison in this work.

In addition, there are a few other methods that are more focused on different aspects of the problem such as enabling joint analysis of multiple datasets (Zeng *et al.* 2013) and incorporating the lineage information between cell types (Biesinger *et al.* 2013). As described in the Introduction section, a number of methods used HMMs to model the chromatin states. These methods differ mainly in the way they model the histone mark data. For instance, some methods discretize the data while others fit Gaussian distributions to the data. In this work we do not explore different methods of modeling the raw histone mark data but rather focus on the parameter estimation technique.

### 2.2 Description of the Hidden Markov Model (HMM)

We use HMMs to represent the chromatin states as hidden states and all possible combinations of (binarized) histone marks as observations. The whole genome is divided into segments of size 200 bp following (Ernst and Kellis 2010, 2012). We define the HMM in matrix form as follows.

Let K be the number of hidden chromatin states and N be the number of possible combinations of histone marks (i.e., *N* = 2*^M^* where *M* = number of histone marks). Let *A* be the state transition matrix where *A_i,j_* is the probability of transition from state *j* to state *i* for 1 *≤ i, j ≤ K*. Let *O* be the emission matrix where *O_i,j_* is the probability of observing the *i*-th combination of the histone marks in state *j* where 1 *≤ i ≤ N* and 1 *≤ j ≤ K*. Let *π* be the initial state distribution vector where *π_i_* is the probability of state *i* in the first segment of each chromosome when 1 *≤ i ≤ K*. For simplicity of the method description, the whole genome is considered as one chromosome by concatenating all chromosomes. Let *T* be the number of segments in the genome. Let *x_t_* be the observation at the *t*-th segment and let *x_t_*_1_:*t*_2_ represent *x_t_*_1_, *x_t_*_1_+1, …, *x_t_*_2_ for *t*_1_ *≤ t*_2_. Given parameters, *θ* = (*A, O, π*), the likelihood of an observed sequence is as follows.

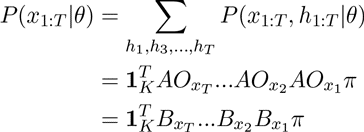

where *h_t_* in the first equation is the hidden state at the *t*-th segment and the summation is taken over all possible sequences of hidden states. *B_i_* is an observable operator (Jaeger 2000) and defined as *B_i_* = *AO_i_*where *O_i_* = *diag*(*O*_*i*,1_, *O_i,_*_2_, …, *O_i,K_)* for 1 *≤ i ≤ N*, and 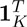 is a vector [1, 1, …, 1] of size *K*.

#### 2.2.1 Inference of the HMM parameters

The method we used was adapted from a method of Mossel and Roch for inferring general phylogenetic tree models (Mossel and Roch 2006). For the observed triple data, (*x_t_, x_t_*_+1_, *x_t_*_+2_), 1 *≤ t ≤ T* – 2, the marginal probabilities of observing the counts of singletons, pairs and triples in the data (the moments in the name “Method of Moments”) are defined as follows:

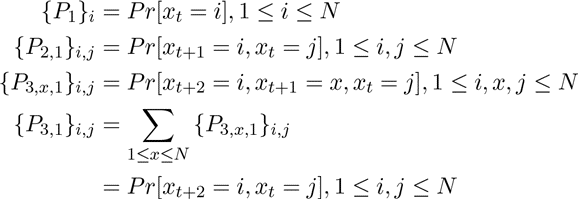

Note that the counts of triples (the third moment) is actually a third-order tensor but for computational reasons we represent it by a collection of matrices indexed by the middle observation *x* where each matrix corresponds to a slice through the tensor. Hsu *et al.* 2012 showed that we can infer the HMM parameters from these marginal probabilities as follows. Let *U* be an *N × K* matrix of the top *K* left singular vectors (computed by the singular value decomposition) of *P*_3_,_1_. Intuitively, *U* acts as surrogate for the observation matrix *O*.

We computed the following matrix *C_x_* for each observation *x* by describing the sample moments using the parameters, *P*_3_*_,x,_*_1_ = *OAO_x_Adiag*(*π*)*O^T^*and *P*_3_,_1_ = *OAAdiag*(*π*)*O^T^*,

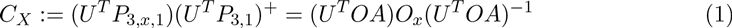

where *M* ^+^ is the Moore-Penrose pseudoinverse of *M*. Since *O_x_* is a diagonal matrix, it is easily seen that *U ^T^ OA* represents the eigenvectors of the matrix and the diagonal elements of *O_x_* are exactly the eigenvalues. We will discuss how to compute the eigenvectors in our practical implementation in section 2.2.2. For now, suppose that the eigenvectors, *U ^T^ OA*, are given. Then for each observation *x*,

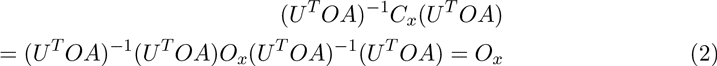

Thus we can infer the emission matrix elements for *x*. The emission matrix *O* is inferred by combining all the *O_x_*’s.

Note that we cannot infer the emission matrix elements directly from the eigenvalues computed from the above matrix *C_x_* because we do not know the order of the eigenvalues. We circumvent this problem by utilizing the fact that the eigenvectors are the same for all observations.

Given the emission probabilities, the remaining parameters of the HMM, *π* and *A*, are easily computed by expressing the sample moments in terms of the parameters, *P*_1_ = *Oπ*, *P*_2_,_1_ = *OAdiag*(*π*)*O^T^*, and the assumption that *A, O, diag*(*π*) are rank *K* (i.e. full-rank) as follows:

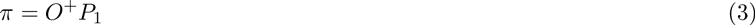

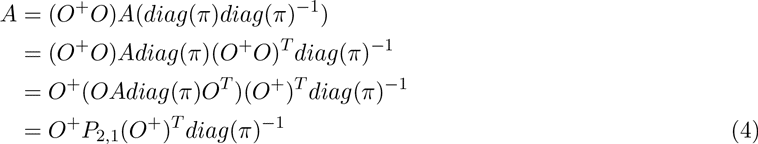

The original algorithm of Hsu *et al.* 2012 assumed that we have access to many short samples from the HMM. When adapting the algorithm to the case of a few long samples, we found that the distribution of initial observations was quite different from the distribution of all observations. Estimating the state initial distribution *π* from the the distribution of all observations *P*_1_ introduces a significant amount of noise. Therefore, we modified *P*_1_ to 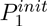 which is the distribution of the first segment of all the chromosomes and we slightly modified Equation 3 as

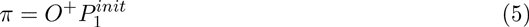

More significantly, we empirically found that most entries of *π* are close to zero and so taking the inverse of *π* in the calculation of *A* would introduce noise. Thus, we modifed the computation of *A* to remove the dependence on *π* as follows:

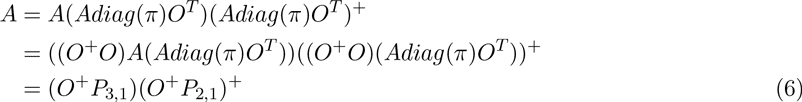

#### 2.2.2 Computing the eigenvectors using anchor observations

It is not easy to compute the eigenvectors correctly because of sample noise. If we compute *U ^T^ OA* from the matrix for each observation, we will generally get different *U ^T^ OA*’s for different observations in practice. Thus, we used a method based on our empirical observations about the histone mark matrix.

The observation space is *N* = 2*^M^* where *M* is the number of histone marks. The maximum number of states is *N*. But, in fact, we observed that *K* is much lower than *N*, consistent with previous papers. Also, for each state, we observed there is a major observation that most of the segments in the state emit. This observation is related to a similar condition considered previously in the topic models setting (Arora *et al.* 2012). For each state *i*, we picked an “anchor observation” 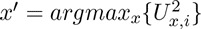 which is similar to using a key word to represent a topic in the topic models setting (Anandkumar *et al.* 2012). The “anchor observation” tends to be the maximum value in the row of *U*, that is, it tends to appear in the state much more often than in the other states. Then we computed eigenvectors from *C_x′_* and extracted a single eigenvector corresponding to the largest eigenvalue, which in practice was separated well from the other eigenvalues. We did this for all states separately and combined the eigenvectors into a single matrix.

##### Algorithm 1: Spectral learning algorithm

**Data**: K: number of states, N: number of observations, T: number of segments Estimate *P*_1_, *P*_2_,_1_, *P*_3_,_1_, and *P*_3_*_,x,_*_1_ from the data.

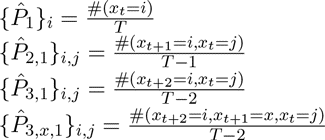

Compute the SVD (singular value decomposition) of 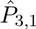 and let *U* be the matrix of the left singular vectors corresponding to the *K* largest singular values.

For each state, compute the matrix *C_x′_* defined in (1) for the anchor observation *x′* and compute the eigenvector *v* corresponding to the largest eigenvalue.

Combine all the eigenvectors *v*’s from the different states.

Infer the emission matrix *O* using (2).

Infer the initial state vector *π* and the transition matrix *A* using (5) and (6).

#### 2.3 Handling technical issues

Since the observation data is noisy, some numerical issues can occur when computing the SVD and eigendecomposition. Previously, (Cohen *et al.* 2013) described implementation issues using spectral learning for another latent variable model in natural language processing. We implemented several optimizations to solve the analogous issues for HMM.

#### 2.3.1 Smoothing of observation matrices

The observation pairs are smoothed out as follows. When *λ* is the smoothing factor between 0 and 1,

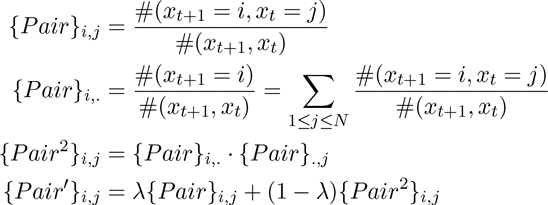

*P air*′ is used as the smoothed observation pair data.

The observation triples are smoothed out using an obvious generalization of the method for smoothing the pairs matrix (see Supplementary Material for details). After the data smoothing, the observation triple and pair matrices are normalized such that the sum of all elements is equal to 1. Intuitively, the smoothing method is similar to adding pseudocounts to sparse data matrices except it uses the marginal frequencies of the observations instead of a uniform pseudocount.

The second and third moments, 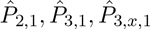 are computed using these smoothed triples and pairs matrices. Specifically, 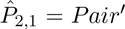 and 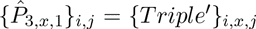 for 1 *≤ i, x, j ≤ N*. We observed that *λ* between 0.9 and 1 seemed to work well for different numbers of states and different cell types. We set *λ* = 0.95 as a default but this parameter is adjustable by the user on the command line of our software. Note that with this setting of *λ* the vast majority of the signal comes from the original data itself and not from the smoothing method.

#### 2.3.2 Handling negative probabilities

Our estimated probabilities should be non-negative but the estimated parameters can have negative values because the signs can be flipped while performing the matrix computations. Therefore, we took the absolute value of the estimated parameters and normalized the parameters following (Cohen *et al.* 2013).

#### 2.3.3 Parameter adjustment

Although we estimated the HMM parameters using spectral learning (1), these estimates are affected by noise in the data and numerical precision errors. Therefore we took the estimates from spectral learning and performed a few iterations of local optimization of the likelihood using EM, following other researchers in the field (K. Chaudhuri, personal communication). The number of iterations was set to five for the further analysis. We call this method **Spectacle** (SPECTral learning for Annotating Chromatin Labels and Epigenomics).

## 3 Results

### 3.1 Data sets and experimental settings

For the histone modification data, we downloaded data for eight histone marks (H3K4me1/2/3, H3K9ac, H3K27ac, H3K27me3, H3K36me3 and H4K20me1) for an ENCODE Tier 1 cell type, GM12878 (version 1) (ENCODE Project Consortium 2012). For external biological validation of our predictions, we used transcription start site (TSS) data using Cap Analysis of Gene Expression (CAGE) from GENCODE (Harrow *et al.* 2012, version 10, May 2012), RNA Polymerase2 (Pol2), P300 and CTCF ChIP-seq data, and polyA RNA-seq data from ENCODE, long intergenic non-coding RNA (lincRNA) RNA-seq data from (Kelley and Rinn 2012), and conserved enhancer regions from the VISTA enhancer browser (Visel *et al.* 2007). All of the data sets except the enhancer data are specific to the cell type that we used for training the HMM.

In order to test the performance of **Spectacle** over multiple cell types, we compared it to **ChromHMM** on data from nine other cell types (H1-hESC, HeLa-S3, HepG2, HMEC, HSMM, HUVEC, K562, NHEK and NHLF).

There are 15,181,508 segments when the human reference genome is divided into 200 bp segments. The histone modification data was binarized by the program, Scripture (Guttman *et al.* (2010)), which was originally developed for transcriptome reconstruction from RNA-seq reads and can also be used for ChIP-seq peak calling. From this, our final data set consisted of presence/absence calls for each histone mark for each segment. Thus the observation space consisted of all combinations of histone marks and in our case the number of observations was 2^8^ = 256. We fixed the number of states to 15 unless stated otherwise, similar to the number of states used in previous studies (Ernst *et al.* 2011). This number of states is readily interpretable biologically and allows us to compare our annotations with previously published annotations.

### 3.2 Spectral learning outperforms the EM algorithm

#### 3.2.1 Initialization of the EM algorithm

The performance of the EM algorithm varied greatly depending on the parameter initialization. For instance, when the number of states was 15, we ran the EM algorithm to convergence for 10 random parameter initializations. The log-likelihood varied from *−*8.33*E*6 to *−*1.11*E*7 (average: *−*9.51*E*6, std. dev.: 7.67*E*5) and the number of iterations needed for the likelihood to converge varied from 16 to 200. Even the highest log-likelihood was lower than that of the heuristic initialization method, *information*, which was *−*7.81*E*6. These results demonstrate the importance of good parameter initializations when running the EM algorithm.

#### 3.2.2 Comparison of the likelihoods

We compared **Spectacle** with **ChromHMM** in terms of the likelihood of the observed data given the parameters. For almost all numbers of states we tested, **Spectacle** had a higher likelihood than **ChromHMM**. The exceptions were when the number of states was 25, 45 and 50 (Table 1). When the number of states was 100, the difference in the performance of the two methods was highly significant and the likelihood found by **ChromHMM** was lower than for smaller numbers of states, which suggests that the EM algorithm found a local optimum at this particular number of states. **ChromHMM** took 20-200 EM iterations to converge to a local optimum in the likelihood. When we ran **ChromHMM** for only five iterations to make its runtime comparable to **Spectacle**, we found that **Spectacle** had a higher likelihood than **ChromHMM** for all numbers of states that we tested. Together these results show that when the runtimes are similar, spectral learning outperforms EM, and even when spectral learning takes a much shorter runtime, it still generally performs better than EM as judged by the likelihood.

**Table 1:** The log-likelihood of **Spectacle** was higher than that of **ChromHMM** as the number of states was varied. Within the parentheses is the log-likelihood ratio between **ChromHMM** with only five EM iterations and **Spectacle**.

| Num of states | ChromHMM | Spectacle | Log-likelihood ratio ($\times 10^4$) |
| --- | --- | --- | --- |
| 5 | -1.74e+07 | **-1.65e+07** | 82.9 |
| 10 | -9.32e+06 | **-9.03e+06** | 29.5 |
| 15 | -7.81e+06 | **-7.17e+06** | 64.2 |
| 20 | -6.31e+06 | **-6.29e+06** | 2.1 |
| 25 | **-5.67e+06** | -5.71e+06 | -3.8 (6.8) |
| 30 | -5.29e+06 | **-5.26e+06** | 2.6 |
| 35 | -5.06e+06 | **-4.93e+06** | 12.9 |
| 40 | -4.75e+06 | **-4.67e+06** | 7.7 |
| 45 | **-4.46e+06** | -4.47e+06 | -1.3 (4.2) |
| 50 | **-4.24e+06** | -4.32e+06 | -8.0 (4.1) |
| 55 | -4.09e+06 | **-4.05e+06** | 3.8 |
| 60 | -3.931e+06 | **-3.87e+06** | 5.9 |
| 65 | -3.80e+06 | **-3.75e+06** | 5.0 |
| 70 | -3.70e+06 | **-3.64e+06** | 6.0 |
| 75 | -3.60e+06 | **-3.56e+06** | 3.9 |
| 80 | -3.53e+06 | **-3.50e+06** | 2.5 |
| 85 | -3.45e+06 | **-3.42e+06** | 3.1 |
| 90 | -3.40e+06 | **-3.36e+06** | 4.5 |
| 95 | -3.34e+06 | **-3.29e+06** | 5.2 |
| 100 | -4.58e+06 | **-3.23e+06** | 134.7 |
| 105 | -3.23e+06 | **-3.18e+06** | 4.9 |

#### 3.2.3 Comparison of training times

We found that **Spectacle** had a significantly faster training time than **ChromHMM** (Figure 1). Spectral learning takes much less compute time than even one iteration of EM and **Spectacle** only needed five iterations of local optimization of the likelihood. Over the range of states that we investigated, **Spectacle** was 3.4-66.8 times (in CPU time) and 4-40 times (in number of iterations) faster than **ChromHMM**.

**Figure 1:**
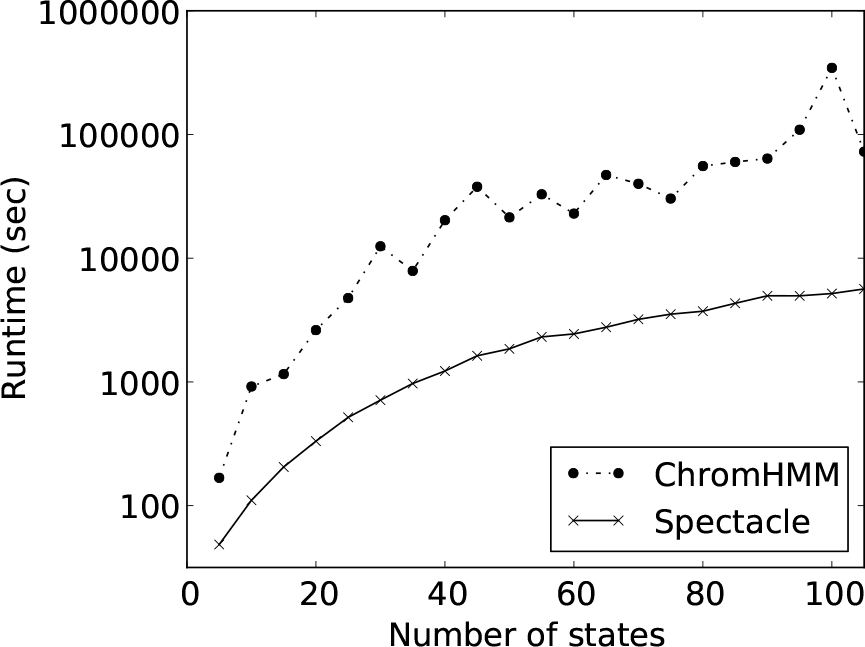
The training time of **Spectacle** vs. **ChromHMM**.

The training time of one iteration of EM is linear in the number of segments and the number of observations, and quadratic in the number of states. On the other hand, the training time of spectral learning is dominated by the computation time of the SVD which is cubic in the number of observations if all singular values are computed exactly (this is the running time of a practical implementation - theoretically the cubic dependence can be improved to the exponent of matrix multiplication (Demmel *et al.* 2007) which is currently *<* 2.373 (Williams 2012)). Thus after the initial creation of the sample moment matrices, the training time of spectral learning is independent of the genome size and can be much faster than the EM algorithm.

#### 3.2.4 Comparison of likelihoods over multiple cell types

We compared both methods over multiple cell types to assess the performance of **Spectacle** over a range of data sets. We examined ten cell types including GM12878 and ran **Spectacle** and **ChromHMM** with 15 hidden chromatin states. **Spectacle** found higher likelihoods for 8 out of 10 cell types while also achieving a 2.6-12.6 fold (in CPU time) and 2.6-13.4 fold (in number of iterations) faster training time (Table 2).

**Table 2:**
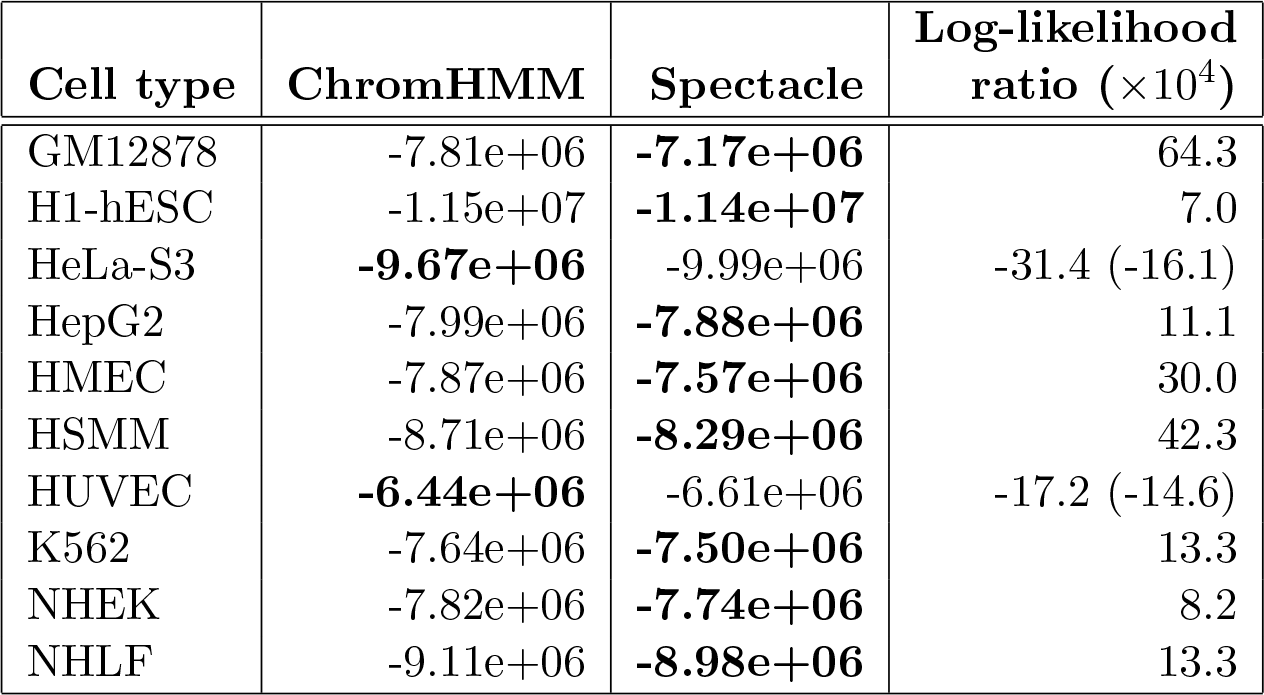
The log-likelihood of **Spectacle** is higher than that of **ChromHMM** for ten cell types. Within the parentheses is the log-likelihood ratio between **ChromHMM** with only five iterations of EM and **Spectacle**.

#### 3.2.5 Empirical sample complexity of the algorithm

Theoretically, the spectral learning algorithm for HMMs is known to have polynomial sample complexity (Hsu *et al.* 2012). To complement these theoretical results, we empirically estimated the sample size required for training the parameters accurately. We trained the parameters of **Spectacle** for 15 states using the whole genome, half the genome, a quarter of the genome, and each chromosome individually using the cell type K562. We then tested the parameters by computing the log-likelihood of the whole genome of another cell type, GM12878, so as not to mix training and testing data sets. We found that the log-likelihood decreased as the sample size decreased as follows: *−*7.62*E*6 for the whole genome; average: *−*7.61*E*6 and std. dev.: 3.6*E*4 for half the genome; average: *−*7.67*E*6 and std. dev.: 1.3*E*5 for a quarter of the genome; and average: *−*8.13*E*6 and std. dev.: 5.7*E*5 for individual chromosomes. This analysis suggests that sample sizes of roughly one chromosome are insufficient for the spectral learning method to accurately train the parameters.

### 3.3 Comparison of the chromatin state annotations

We estimated the HMM parameters using **Spectacle** for 15 chromatin states for one of the ENCODE cell lines, GM12878. Then we assigned the hidden states to segments using the posterior decoding algorithm, which assigns the most likely state to each segment. We used this method for both **Spectacle** and EM. We performed external validation of the predicted states using experimental data other than the histone modification data. We computed the enrichment of signals such as histone modification marks and independent biological datasets in the hidden HMM states (Figure S1,S2). The hidden states were manually annotated as TSS (transcription start site), enhancer, lincRNA, or exon (transcribed regions) according to the enrichment of signals using a similar procedure to previous works (Ernst and Kellis 2010; Hoffman *et al.* 2013) (see Supplementary Material for details).

**Figure S1:**
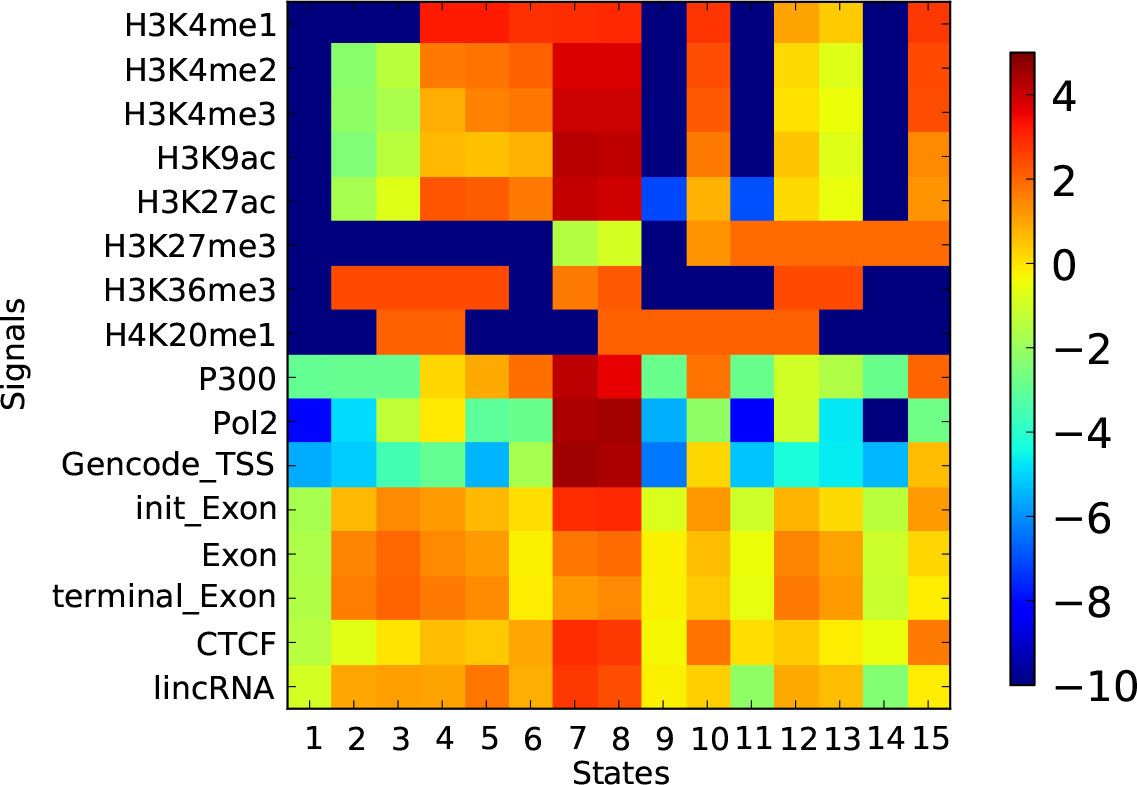
The enrichment of signals for each state when the number of states is 15 for **Spectacle**. TSS states are 7 and 8, enhancer states are 4, 10, 12 and 15, lincRNA states are 3, 5, 7 and 8, and transcribed exon states are 2, 3, 4, 5, 7, 8, 12 and 13.

**Figure S2:**
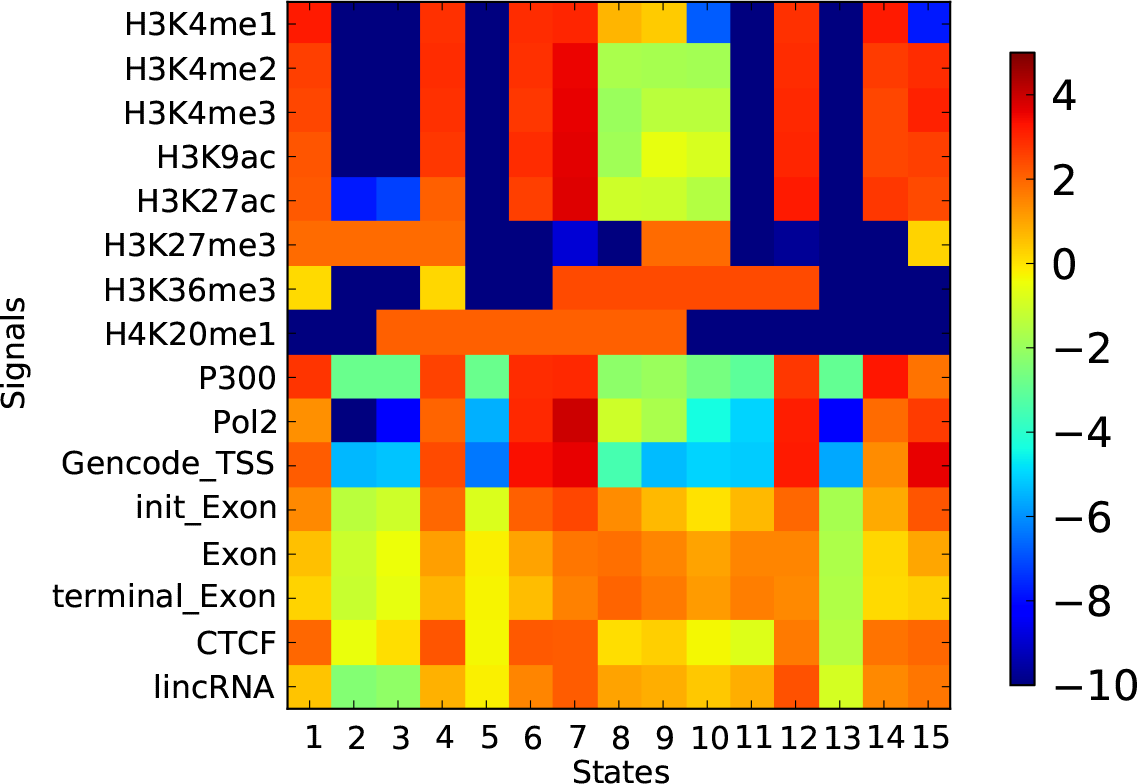
The enrichment of signals for each state when the number of states is 15 for **ChromHMM**. TSS states are 4, 7, 12 and 15, enhancer states are 1, 6, 9 and 14, lincRNA states are 7, 8, 12 and 15, and transcribed exon states are 1, 4, 7, 8, 9, 10, 11 and 12.

Two states for **Spectacle** and four states for **ChromHMM** were annotated as TSS. **Spectacle** predicted TSS or TSS plus flanking regions(*±*1 kb) significantly more accurately than **ChromHMM** while using a smaller number of states (Figure 2,3). TSS regions represent only 0.07% of the whole genome. About 93% of TSSs were found within the inferred TSS states in the HMM consisting of about 4% of the genome (*∼*22 fold enrichment). Also, the flanking regions (*±*1 kb) of TSS including promoter regions have similar histone mark patterns to TSS. About 89% of TSS flanking regions was found within the TSS states (*∼*21-fold enrichment). Furthermore, the states have the highest enrichment of H3K4me3, H3K27ac, Pol2 and initial exon among the states, all of which are indicators of TSS and promoters (Figures S1,S2).

**Figure 2:**
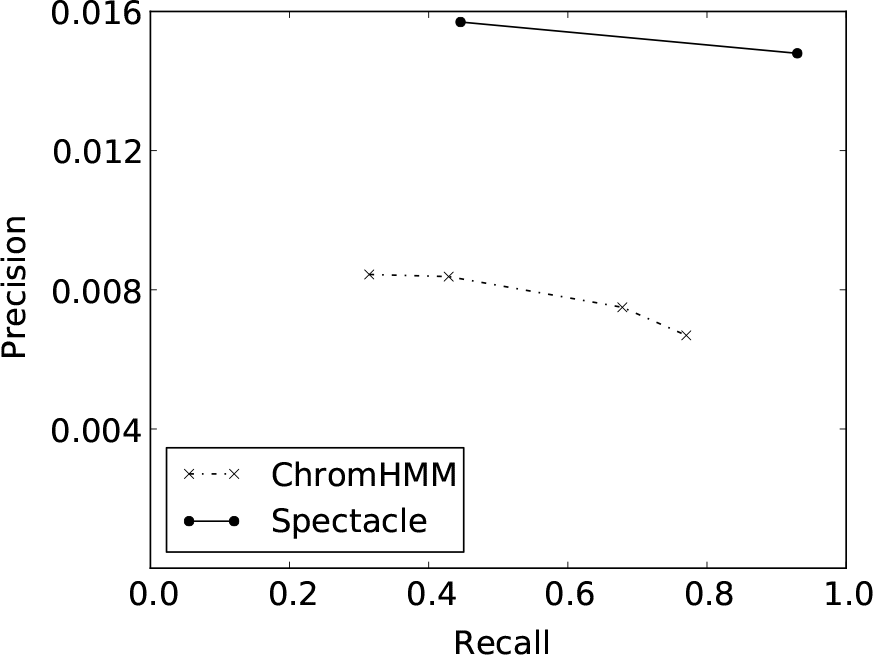
Precision-Recall curve for prediction of TSS when the number of states is 15.

**Figure 3:**
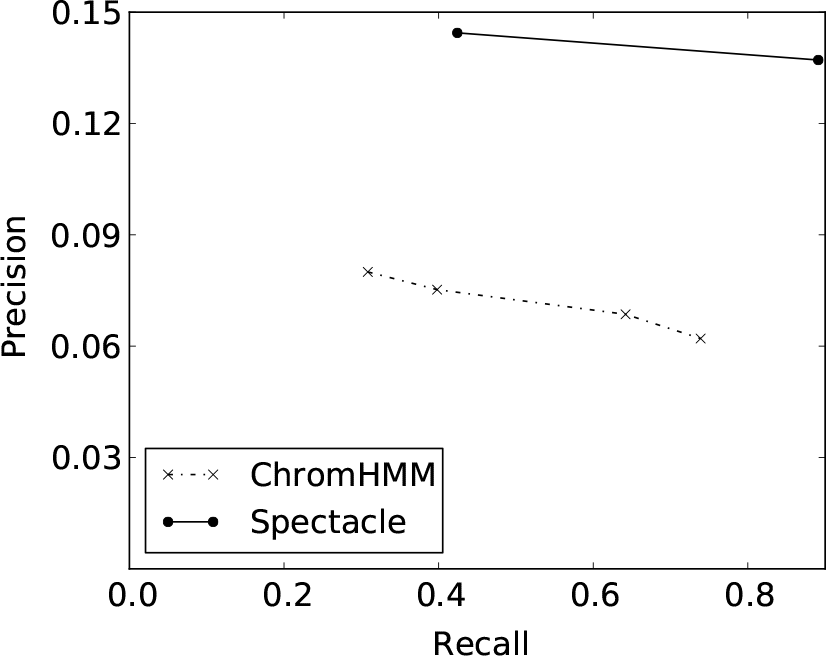
Precision-Recall curve for prediction of TSS plus flanking regions (within *±* 1 kb) when the number of states is 15.

Transcribed regions also have specific epigenetic patterns (Figure 4). The TSS states and six other states for **Spectacle** and three out of the four TSS states and five other states for **ChromHMM** were annotated as Exon states (*∼*3-fold enrichment), and they were enriched with H3K36me3, which is a signal for gene bodies (Bannister *et al.* 2005).

**Figure 4:**
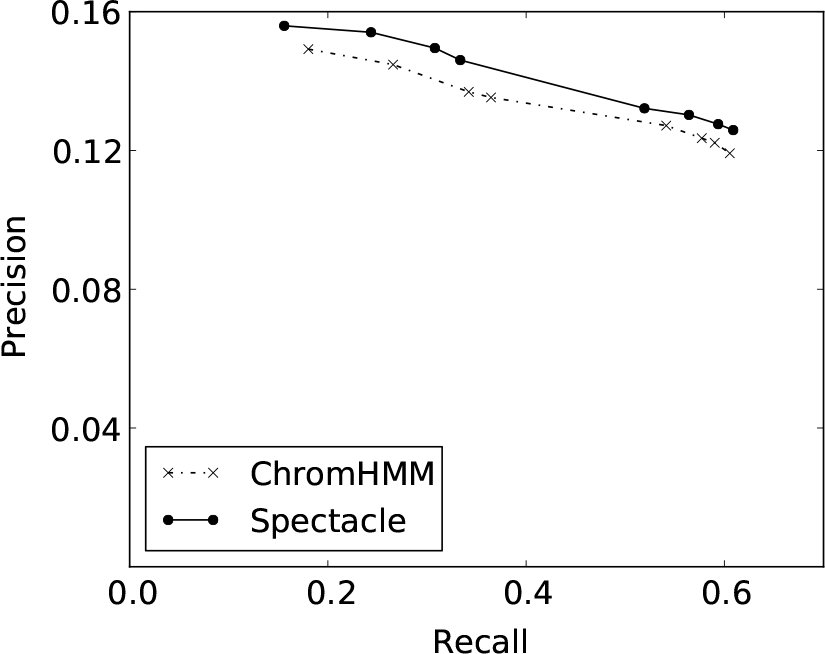
Precision-Recall curve for prediction of the transcribed exons when the number of states is 15.

Another important class of functional elements is lincRNAs (long intergenic non-coding RNAs) (Figure 5). It is known that lincRNAs have the “K4-K36 signature” (Khalil *et al.* 2009), that is, H3K4me3 enrichment at the transcription start site and H3K36me3 throughout the transcribed regions. Since this histone pattern is similar to protein-coding genes, lincRNAs are identified by searching for K4-K36 domains outside of known protein-coding genes. In fact, the TSS states and two of the Exon states for **Spectacle** and three out of the four TSS states and one Exon state for **ChromHMM** were annotated as lincRNAs (*∼*4-fold enrichment).

**Figure 5:**
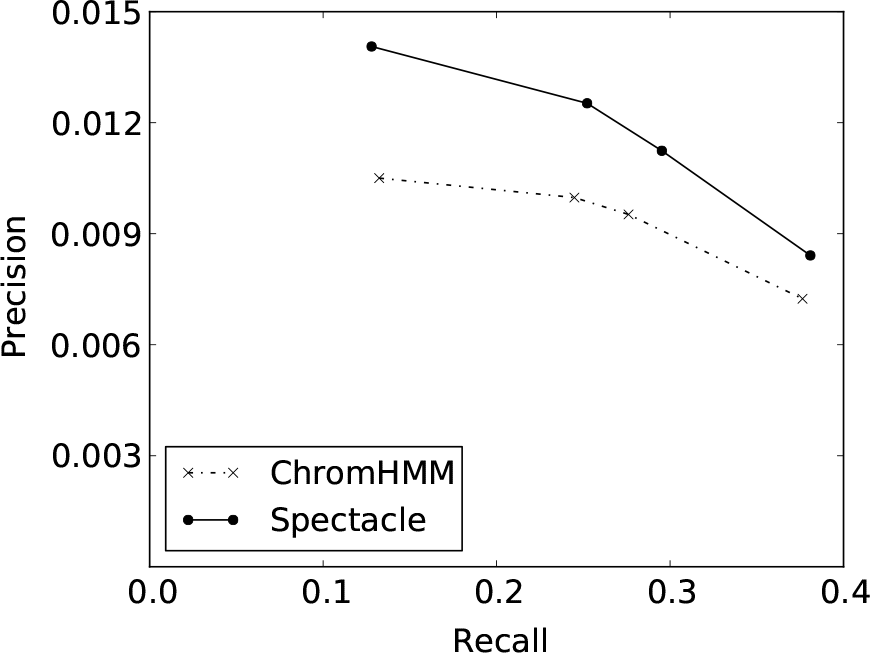
Precision-Recall curve for prediction of lincRNAs when the number of states is 15.

Lastly, four states for both **Spectacle** and **ChromHMM** were annotated as Enhancer (Figure 6) (*∼*2-fold enrichment). The Enhancer states did not overlap with TSS states and had a higher enrichment of H3K4me1/2 than that of H3K4me3. Two states among the Enhancer states had high enrichment of both P300 and CTCF.

**Figure 6:**
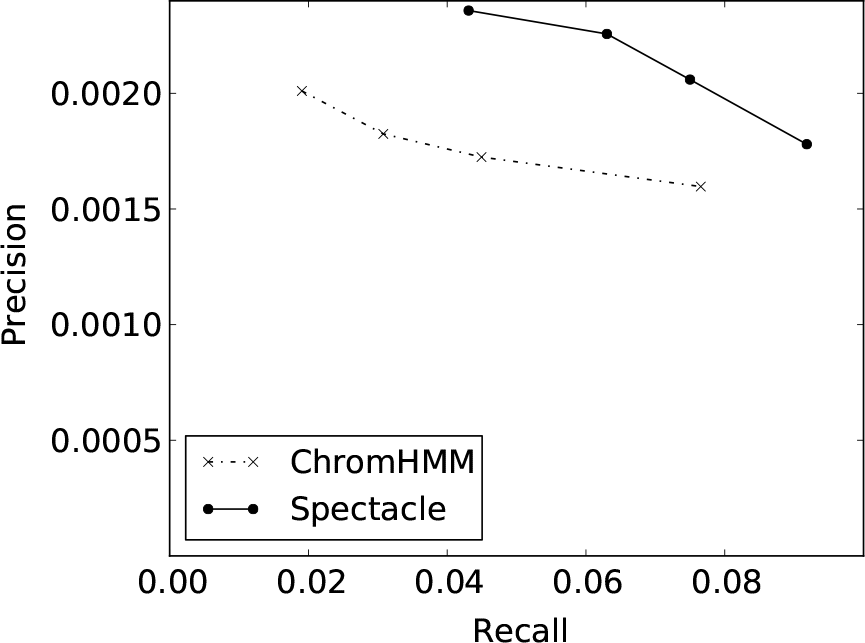
Precision-Recall curve for prediction of enhancers when the number of states is 15.

Overall, our results showed improvements in predicting all classes of functional elements that we tested, compared to a previous state-of-the-art method based on the EM algorithm.

## 4 Discussion

We have presented an improved method and software tool called **Spectacle** (SPECTral learning for Annotating Chromatin Labels and Epigenomics) for annotating chromatin states in the human genome. At the heart of our method is a state-of-the-art spectral learning method for unsupervised learning of the parameters of a Hidden Markov Model (HMM). To implement the spectral learning algorithm in practice, we made several technical modifications which improved its accuracy and numerical stability on the data sets we tested. These modifications may be of broader interest in other applications, including other computational biology problems, computer vision and natural language processing. We showed that **Spectacle** outperforms a previous state-of-the-art method, ChromHMM, by finding higher likelihoods in most data sets tested, having a much faster training time and more accurately annotating several independent biological datasets. Recently, the ChromHMM code was updated to add an option for learning parameters in parallel using multiprocessors. Using the option would improve the runtime of both **Spectacle** and **ChromHMM** (http://compbio.mit.edu/ChromHMM/). Our software implementation is freely available online and is lightweight and easy to use on a regular desktop without the need for specialized computer hardware. Our code modifies the ChromHMM code which has been used by several groups so we believe it will be user-friendly and accessible.

Spectral learning can be applied to multiple problems in computational biology. A recent work used spectral learning for poly(A) motif prediction (Xie *et al.* 2013). The authors did not try to recover the HMM parameters explicitly but instead learned them up to an unknown invertible linear transformation and used the transformed parameters as features for classification by a Support Vector Machine. Another recent paper (Zou *et al.* 2013) applied a spectral learning algorithm for contrastive learning to a problem involving epigenome maps. This is a more restricted version of the problem we study here in which one is specifically contrasting two data sets instead of annotating a single data set. Since Hidden Markov Models and the Expectation-Maximization algorithm are used in many other problems in computational biology, it is possible that the methods described here may be useful in those settings.

In the future, having a faster and more accurate chromatin state annotation tool should be useful for annotating multiple epigenomic maps. Previous works found associations between disease-causing variants and epigenomic marks (Maurano *et al.* 2012; Kasowski *et al.* 2013; Kilpinen *et al.* 2013; McVicker *et al.* 2013), suggesting that better understanding of the epigenome might help interpret the variants underlying human disease. For example, Kasowski *et al.* 2013 used ChromHMM to infer variation in chromatin states across individuals, so we expect **Spectacle** to be useful for other similar types of data sets in the future. Indeed there are currently several different major epigenomics projects (e.g., BLUEPRINT, Roadmap Epigenomics Project), producing chromatin mark data for many cell types and human population (Adams *et al.* 2012; Bernstein *et al.* 2010). Given the rapid decrease in the cost of sequencing, we also expect that many more epigenomic maps will be produced in different cell types, human populations (Kasowski *et al.* 2013), species, environmental conditions and developmental contexts (Ernst *et al.* 2011; Zhu *et al.* 2013). Thus we expect that the need for fast and accurate tools for processing this type of data will continue to grow in the future.

## 5 Acknowledgements

We thank Kamalika Chaudhuri, Alexander Schliep and Mona Singh for helpful comments and discussions. We also thank Jason Ernst for making the ChromHMM code freely available online and for helpful comments on a previous version of this manuscript. This work was partially funded by the National Institutes of Health (R00HG004515 to Kevin C. Chen).

## 6 Supplement

### Parameter initialization method: “information”

For completeness we briefly describe the parameter initialization method described by Ernst and Kellis. We refer the reader to the original paper for a full description. The number of hidden states is denoted as K. The “information” method iteratively partitions the segments of the genome into K groups based on the presence of a selected chromatin mark. The group assignment for all segments is considered as a crude prediction of the hidden states, and the emission, transition and initial probability parameters are inferred accordingly from the hidden state prediction.

The “information” method assigns groups to segments as follows. In the first step, all segments are assigned to group 1. Then for each step, the method creates a new group by partitioning an existing group into two groups. It repeats this partitioning step (K-1) times. To choose a group to partition, the method finds the best group and chromatin mark for splitting the group that maximizes an entropy measure. The segments in the group which have the mark constitute a new group and the segments in the group which do not have the mark remain in the group.

## Data smoothing for triples

The observation triples (*x_t_, x_t_*_+1_, *x_t_*_+2_) are smoothed out as follows.

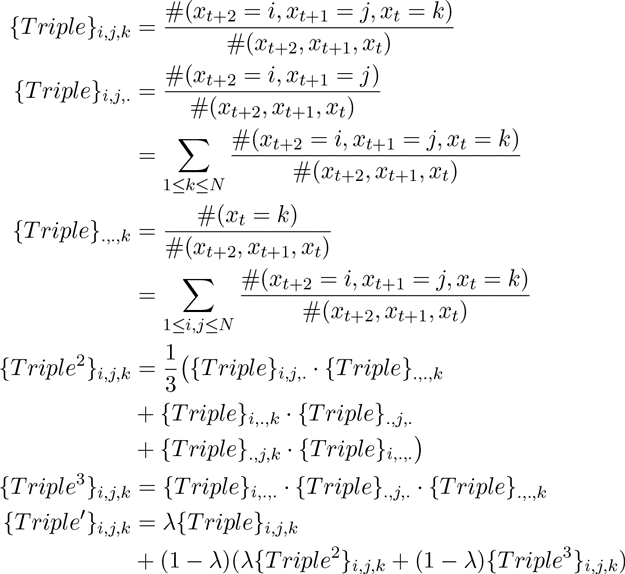

*Triple*′ is used as the smoothed observation triple data.

### Enrichment of biological signals and state annotation

The enrichment of a signal (either a chromatin modification or a biological annotation) for a state is calculated as follows:

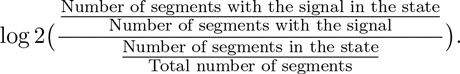

Our approach is semi-automated in the sense that we assign biological interpretations to the hidden states manually, similar to previous works (Ernst and Kellis (2010); Hoffman *et al.* (2013)). We based our manual annotations on experimental evidence from (Barski *et al.* (2007); Heintzman *et al.* (2009)). States with positive enrichment of Pol2, H3K27ac, H3K4me3 and H3K3me1, as well as greater enrichment of H3K4me3 compared to H3K4me1 were annotated as “TSS”. States with positive enrichment of Enhancer, H3K4me3 and H3K3me1, as well as greater enrichment of H3K4me1 compared to H3K4me3 were annotated as “Enhancer”. States with positive enrichment of lincRNA and positive enrichment of either H3K4me3 or H3K36me3 but not annotated as “Enhancer” were annotated as “lincRNA”. States with positive enrichment of both Exon and H3K36me3 were annotated as “Exon”.

Instead of the above manual assignments, we also ranked states by enrichment for each biological dataset and plotted the Precision-Recall curves. The results were consistent in that Spectacle always performed the same or better than ChromHMM.

## References

Adams, D., Altucci, L., Antonarakis, S. E., Ballesteros, J., Beck, S., Bird, A., Bock, C., Boehm, B., Campo, E., Caricasole, A., Dahl, F., Dermitzakis, E. T., Enver, T., Esteller, M., Estivill, X., Ferguson-Smith, A., Fitzgibbon, J., Flicek, P., Giehl, C., Graf, T., Grosveld, F., Guigo, R., Gut, I., Helin, K., Jarvius, J., Küppers, R., Lehrach, H., Lengauer, T., Åke Lernmark, Leslie, D., Loeffler, M., Macintyre, E., Mai, A., Martens, J. H., Minucci, S., Ouwehand, W. H., Pelicci, P. G., Pendeville, H., Porse, B., Rakyan, V., Reik, W., Schrappe, M., Schübeler, D., Seifert, M., Siebert, R., Simmons, D., Soranzo, N., Spicuglia, S., Stratton, M., Stunnenberg, H. G., Tanay, A., Torrents, D., Valencia, A., Vellenga, E., Vingron, M., Walter, J., and Willcocks, S. (2012). BLUEPRINT to decode the epigenetic signature written in blood. Nature Biotechnology, 30, 224–226.

Anandkumar, A., Hsu, D., and Kakade, S. M. (2012). A method of moments for mixture models and hidden Markov models. In Proceedings of the 25th Conference on Learning Theory (COLT).

Arora, S., Ge, R., and Moitra, A. (2012). Learning topic models – Going beyond SVD. In IEEE 53rd Annual Symposium on Foundations of Computer Science (FOCS).

Bannister, A. J., Schneider, R., Myers, F. A., Thorne, A. W., Crane-Robinson, C., and Kouzarides, T. (2005). Spatial distribution of Di- and Tri-methyl Lysine 36 of Histone H3 at active genes. Journal of Biological Chemistry, 280(18), 17732–17736.

Barski, A., Cuddapah, S., Cui, K., Roh, T.-Y., Schones, D. E., Wang, Z., Wei, G., Chepelev, I., and Zhao, K. (2007). High-resolution profiling of histone methylations in the human genome. Cell, 129, 823–837.

Bernstein, B. E., Stamatoyannopoulos, J. A., Costello, J. F., Ren, B., Milosavljevic, A., Meissner, A., Kellis, M., Marra, M. A., Beaudet, A. L., Ecker, J. R., Farnham, P. J., Hirst, M., Lander, E. S., Mikkelsen, T. S., and Thomson, J. A. (2010). The NIH Roadmap Epigenomics Mapping Consortium. Nature Biotechnology, 28, 1045–1048.

Biesinger, J., Wang, Y., and Xie, X. (2013). Discovering and mapping chromatin states using a tree hidden Markov model. BMC Bioinformatics, 14(Suppl 5), S4.

Cohen, S., Stratos, K., Collins, M., Foster, D., and Ungar, L. (2013). Experiments with spectral learning of latent variable PCFGs. In Preceedings of the 2013 Conference of the North American Chapter of the Association for Computational Linguistics: Human Language Technologies (NAACL).

Demmel, J., Dumitriu, I., and Holtz, O. (2007). Fast linear algebra is stable. Journal Numerische Mathematik, 108(1), 59–91.

Dempster, A., Laird, N., and Rubin, D. (1977). Maximum likelihood from incomplete data via the EM algorithm. J. Roy. Stat. Soc., 39(1), 1–38.

ENCODE Project Consortium (2012). An integrated encyclopedia of DNA elements in the human genome. Nature, 489, 57–74.

Ernst, J. and Kellis, M. (2010). Discovery and characterization of chromatin states for systematic annotation of the human genome. Nature Biotechnology, 28(8), 817–825.

Ernst, J. and Kellis, M. (2012). ChromHMM: automating chromatin state discovery and characterization. Nature Methods, 9(3), 215–216.

Ernst, J., Kheradpour, P., Mikkelsen, T. S., Shoresh, N., Ward, L. D., Epstein, C. B., Zhang, X., Wang, L., Issner, R., Coyne, M., Ku, M., Durham, T., Kellis, M., and Bernstein, B. E. (2011). Mapping and analysis of chromatin state dynamics in nine human cell types. Nature, 473, 43–49.

Filion, G. J., van Bemmel, J. G., Braunschweig, U., Talhout, W., Kind, J., Ward, L. D., Brugman, W., de Castro, I. J., Kerkhoven, R. M., Bussemaker, H. J., and van Steensel, B. (2010). Systematic protein location mapping reveals five principal chromatin types in Drosophila cells. Cell, 143, 212–224.

Guttman, M., Amit, I., Garber, M., French, C., Lin, M. F., Feldser, D., Huarte, M., Zuk, O., Carey, B. W., Cassady, J. P., Cabili, M. N., Jaenisch, R., Mikkelsen, T. S., Jacks, T., Hacohen, N., Bernstein, B. E., Kellis, M., Regev, A., Rinn, J. L., and Lander, E. S. (2009). Chromatin signature reveals over a thousand highly conserved large non-coding RNAs in mammals. Nature, 458, 223–227.

Guttman, M., Garber, M., Levin, J. Z., Donaghey, J., Robinson, J., Adiconis, X., Fan, L., Koziol, M. J., Gnirke, A., Nusbaum, C., Rinn, J. L., Lander, E. S., and Regev, A. (2010). Ab initio reconstruction of cell typespecific transcriptomes in mouse reveals the conserved multi-exonic structure of lincRNAs. Nature Biotechnology, 28(5), 503–510.

Harrow, J., Frankish, A., Gonzalez, J., Tapanari, E., Diekhans, M., Kokocinski, F., Aken, B., Barrell, D., Zadissa, A., Searle, S., Barnes, I., Bignell, A., Boychenko, V., Hunt, T., Kay, M., Mukherjee, G., Rajan, J., Despacio-Reyes, G., Saunders, G., Steward, C., Harte, R., Lin, M., Howald, C., Tanzer, A., Derrien, T., Chrast, J., Walters, N., Balasubramanian, S., Pei, B., Tress, M., Rodriguez, J., Ezkurdia, I., van Baren, J., Brent, M., Haussler, D., Kellis, M., Valencia, A., Reymond, A., Gerstein, M., Guigo, R., and Hubbard, T. (2012). GENCODE: the reference human genome annotation for The ENCODE Project. Genome Research, 22(9), 1760–1774.

Heintzman, N. D., Hon, G. C., Hawkins, R. D., Kheradpour, P., Stark, A., Harp, L. F., Ye, Z., Lee, L. K., Stuart, R. K., Ching, C. W., Ching, K. A., Antosiewicz-Bourget, J. E., Liu, H., Zhang, X., Green, R. D., Lobanenkov, V. V., Stewart, R., Thomson, J. A., Crawford, G. E., Kellis, M., and Ren, B. (2009). Histone modifications at human enhancers reflect global cell-type-specific gene expression. Nature, 459, 108–112.

Hoffman, M. M., Buske, O. J., Wang, J., Weng, Z., Bilmes, J. A., and Noble, W. S. (2012). Unsupervised pattern discovery in human chromatin structure through genomic segmentation. Nature Methods, 9(5), 473–476.

Hoffman, M. M., Ernst, J., Wilder, S. P., Kundaje, A., Harris, R. S., Libbrecht, M., Giardine, B., Ellenbogen, P. M., Bilmes, J. A., Birney, E., Hardison, R. C., Dunham, I., Kellis, M., and Noble, W. S. (2013). Integrative annotation of chromatin elements from ENCODE data. Nucleic Acids Research, 41(2), 827–841.

Hsu, D., Kakade, S., and Zhang, T. (2012). A spectral algorithm for learning hidden Markov models. Journal of Computer and System Sciences, 78, 1460–1480.

Huang, X., Acero, A., and Hon, H.-W. (2001). Spoken language processing. Prentice-Hall, Upper Saddle River, NJ.

Jaeger, H. (2000). Observable operator models for discrete stochastic time series. Neural Computation, 12(6), 1371–1398.

Jaschek, R. and Tanay, A. (2009). Spatial clustering of multivariate genomic and epigenomic information. Research in Computational Molecular Biology (RECOMB), LNCS, 5541, 170–183.

Kasowski, M., Kyriazopoulou-Panagiotopoulou, S., Grubert, F., Judith B. Zaugg,., Kundaje, A., Liu, Y., Boyle, A. P., Zhang, Q. C., Zakharia, F., Spacek, D. V., Li, J., Xie, D., Olarerin-George, A., Steinmetz, L. M., Hogenesch, J. B., Kellis, M., Batzoglou, S., and Snyder, M. (2013). Extensive variation in chromatin states across humans. Science, 342, 750–752.

Kelley, D. and Rinn, J. (2012). Transposable elements reveal a stem cell-specific class of long noncoding RNAs. Genome Biology, 13, R107.

Khalil, A. M., Guttman, M., Huarte, M., Garbera, M., Rajd, A., Morales, D. R., Thomas, K., Presser, A., Bernstein, B. E., van Oudenaarden, A., Regev, A., Lander, E. S., and Rinn, J. L. (2009). Many human large intergenic noncoding RNAs associate with chromain-modifying complexes and affect gene expression. Proceedings of the National Academy of Sciences of the United States of America, 106(28), 11667–11672.

Kharchenko, P. V., Alekseyenko, A. A., Schwartz, Y. B., Minoda, A., Riddle, N. C., Ernst, J., Sabo, P. J., Larschan, E., Gorchakov, A. A., Gu, T., Linder-Basso, D., Plachetka, A., Shanower, G., Tolstorukov, M. Y., Luquette, L. J., Xi, R., Jung, Y. L., Park, R. W., Bishop, E. P., Canfield, T. K., Sandstrom, R., Thurman, R. E., MacAlpine, D. M., Stamatoyannopoulos, J. A., Kellis, M., Elgin, S. C. R., Kuroda, M. I., Pirrotta, V., Karpen, G. H., and Park, P. J. (2011). Comprehensive analysis of the chromatin landscape in Drosophila melanogaster. Nature, 471, 480–485.

Kilpinen, H., Waszak, S. M., Gschwind, A. R., Raghav, S. K., Witwicki, R. M., Orioli, A., Migliavacca, E., Wiederkehr, M., Gutierrez-Arcelus, M., Panousis, N. I., Yurovsky, A., Lappalainen, T., Romano-Palumbo, L., Planchon, A., Bielser, D., Bryois, J., Padioleau, I., Udin, G., Thurnheer, S., Hacker, D., Core, L. J., Lis, J. T., Hernandez, N., Reymond, A., Deplancke, B., and Dermitzakis, E. T. (2013). Coordinated effects of sequence variation on DNA binding, chromatin structure, and transcription. Science, 342, 744–747.

Lai, W. K. M. and Buck, M. J. (2013). An integrative approach to understanding the combinatorial histone code at functional elements. Bioinformatics, 29(18), 2231–2237.

Lian, H., Thompson, W. A., Thurman, R., Stamatoyannopoulos, J. A., Noble, W. S., and Lawrence, C. E. (2008). Automated mapping of large-scale chromatin structure in ENCODE. Bioinformatics, 24, 1911–1916.

Maurano, M. T., Humbert, R., Rynes, E., Thurman, R. E., Haugen, E., Wang, H., Reynolds, A. P., Sandstrom, R., Qu, H., Brody, J., Shafer, A., Neri, F., Lee, K., Kutyavin, T., Stehling-Sun, S., Johnson, A. K., Canfield, T. K., Giste, E., Diege, M., Bates, D., Hansen, R. S., Neph, S., Sabo, P. J., Heimfeld, S., Raubitschek, A., Ziegler, S., Cotsapas, C., Sotoodehnia, N., Glass, I., Sunyaev, S. R., Kaul, R., and Stamatoyannopoulos, J. A. (2012). Systematic localization of common disease-associated variation in regulatory DNA. Science, 337, 1190–1195.

McVicker, G., van de Geijn, B., Degner, J. F., Cain, C. E., Banovich, N. E., Raj, A., Lewellen, N., Myrthil, M., Gilad, Y., and Pritchard, J. K. (2013). Identification of genetic variants that affect histone modifications in human cells. Science, 342, 747–749.

Mossel, E. and Roch, S. (2006). Learning nonsingular phylogenies and hidden Markov models. The Annals of Applied Probability, 16(2), 583–614.

Pearson, K. (1895). Contributions to the mathematical theory of evolution. Philosophical Transactions of the Royal Society of London, A, 186, 343–414.

Portela, A. and Esteller, M. (2010). Epigenetic modifications and human disease. Nature Biotechnology, 26(10), 1057–1068.

Rabiner, L. R. (1989). A tutorial on hidden Markov models and selected applications in speech recognition. Proceedings of the IEEE, 77(2), 257–286.

Rivera, C. M. and Ren, B. (2013). Mapping human epigenomes. Cell, 155(1), 39–55.

Shen, Y., Yue, F., McCleary, D. F., Ye, Z., Edsall, L., Kuan, S., Wagner, U., Dixon, J., Lee, L., Lobanenkov, V. V., and Ren, B. (2012). A map of the cis-regulatory sequences in the mouse genome. Nature, 488, 116–120.

Ucar, D., Hu, Q., and Tan, K. (2011). Combinatorial chromatin modification patterns in the human genome revealed by subspace clustering. Nucleic Acids Research, 39(10), 4063–4075.

Visel, A., Minovitsky, S., Dubchak, I., and Pennacchio, L. A. (2007). VISTA Enhancer Browser–a database of tissue-specific human enhancers. Nucleic Acids Research, 35**(Database issue)**, D88–D92.

Wang, J., Lunyak, V. V., and Jordan, I. K. (2012). Chromatin signature discovery via histone modification profile alignments. Nucleic Acids Research, 40(21), 10642–10656.

Williams, V. V. (2012). Multiplying matrices faster than Coppersmith-Winograd. In Symposium on the Theory of Computing (STOC).

Won, K.-J., Zhang, X., Wang, T., Ding, B., Raha, D., Snyder, M., Ren, B., and Wang, W. (2013). Comparative annotation of functional regions in the human genome using epigenomic data. Nucleic Acids Research, 41(8), 4423–4432.

Xie, B., Jankovic, B., Bajic, V., Song, L., and Gao, X. (2013). Poly(A) motif prediction using spectral latent features from human DNA sequences. Bioinformatics, 29(13), i316–i325.

Yip, K. Y., Cheng, C., Bhardwaj, N., Brown, J. B., Leng, J., Kundaje, A., Rozowsky, J., Birney, E., Bickel, P., Snyder, M., and Gerstein, M. (2012). Classification of human genomic regions based on experimentally determined binding sites of more than 100 transcription-related factors. Genome Biolgoy, 13, R48.

Zeng, X., Sanalkumar, R., Bresnick, E. H., Li, H., Chang, Q., and Keles, S. (2013). jMO-SAiCS: joint analysis of multiple ChIP-seq datasets. Genome Biology, 14, R38.

Zhu, J., Adli, M., Zou, J. Y., Verstappen, G., Coyne, M., Zhang, X., Durham, T., Miri, M., Deshpande, V., Jager, P. L. D., Bennett, D. A., Houmard, J. A., Muoio, D. M., Onder, T. T., Camahort, R., Cowan, C. A., Meissner, A., Epstein, C. B., Shoresh, N., and Bernstein, B. E. (2013). Genome-wide chromatin state transitions associated with developmental and environmental cues. Cell, 152, 642–654.

Zou, J., Hsu, D., Parkes, D., and Adams, R. (2013). Contrastive learning using spectral methods. In Advances in Neural Information Proceeding Systems (NIPS).

